# A novel zebrafish-based *in vivo* model of Zika virus infection unveils NS4A as a key viral determinant of neuropathogenesis

**DOI:** 10.1101/2024.01.20.576493

**Authors:** Aïssatou Aïcha Sow, Priyanka Jamadagni, Pietro Scaturro, Shunmoogum A. Patten, Laurent Chatel-Chaix

## Abstract

Infection of pregnant women by Zika virus (ZIKV) is associated with severe neurodevelopmental defects in newborns through poorly defined mechanisms. Here, we engineered a zebrafish *in vivo* model of ZIKV infection to circumvent limitations of existing mammalian models. Leveraging the unique tractability of this system, we gained unprecedented access to the ZIKV-infected brain at early developmental stages. The infection of zebrafish larvae with ZIKV phenocopied the disease in mammals including a reduced head area and neural progenitor cells (NPC) infection and depletion. Moreover, transcriptomic analyses of ZIKV-infected NPCs revealed a distinct dysregulation of genes involved in survival and neuronal differentiation, including downregulation of the expression of the glutamate transporter *vglut1*, resulting in an altered glutamatergic network in the brain. Mechanistically, ectopic expression of ZIKV protein NS4A in the larvae recapitulated the morphological defects observed in infected animals, identifying NS4A as a key determinant of neurovirulence and a promising antiviral target for developing therapies.

## INTRODUCTION

Since its introduction and emergence in Latin America in 2015, Zika virus (ZIKV) constitutes a public health concern, especially considering that neither antiviral treatments nor vaccines against this flavivirus are currently available. ZIKV is mainly transmitted through mosquito bite but unlike most flaviviruses, it can also be vertically transmitted to the fetus in infected pregnant women. ZIKV may reach the fetal brain and cause major neurodevelopment defects leading to severe birth defects and life-long disabilities. This affliction named congenital Zika syndrome (CZS) includes microcephaly, brain damage and body mobility restriction. Even asymptomatic infected women (80% of the cases) are at risk of delivering newborns with long-term neurological/cognitive problems.^1^ Notably, children that were exposed to ZIKV *in utero* but without CZS at birth may nevertheless show neurodevelopmental delays within the first two years of life, particularly in cognitive and language development.^2^ These symptoms can be explained by the fact that following congenital transmission, ZIKV infects neural progenitor cells (NPC) in the fetal brain resulting in an alteration of their differentiation program as well as in their apoptosis-driven cell death through poorly defined mechanisms.^3–7^ In addition, ZIKV also infects other cell types of the brain such as astrocytes and microglial cells, potentially inducing neuroinflammation and thereby, indirectly contributing to the brain developmental defects.^8^ Ultimately, brain infection leads to severe defects in neuronal maturation resulting in cortical thinning, growth restriction and thus, reduced size of the brain. Importantly, the viral and host determinants driving ZIKV neuropathogenesis are largely misunderstood.

Upon ZIKV entry into the target cell, the positive-sense viral RNA (vRNA) genome is translated into a polyprotein which is subsequently cleaved into 10 mature viral proteins. The seven non-structural (NS) proteins are responsible for vRNA replication while structural proteins capsid (C), pre-membrane (prM) and envelope (E), together with the viral genome, orchestrate the assembly of new viral particles.^9,10^ Flaviviral NS4A and NS4B are of particular interest since they are absolutely required for replication through poorly defined process and in the case of ZIKV, were reported to inhibit the growth of NPC-derived neurospheres *in vitro* alone or in combination.^11–16^ These transmembrane proteins lack enzymatic activities and interact together in the endoplasmic reticulum (ER) suggesting that they share functions in viral replication. Reverse genetics and pharmacological approaches recently demonstrated that NS4A of ZIKV and dengue virus (DENV, closely related to ZIKV) contribute to the biogenesis of ER-derived replication organelles which host the vRNA synthesis process.^12,17^ In addition, the NS4A-NS4B precursor is also believed to play specific roles in flavivirus life cycle.^18^ Interestingly, DENV NS4B is the target of a several highly potent antivirals, including two currently in phase 2 clinical trials.^19–23^ Furthermore, individual expression of ZIKV NS4A and NS4B can inhibit the AKT/mTOR pathway and interfere with the early induction of type I interferon.^11,24^ Finally, ubiquitous expression of ZIKV NS4A in the invertebrate model of drosophila led to marked decrease in the size of the third instar larval brain^25,26^, although such impact remains to be confirmed in a vertebrate model whose CNS development resembles more closely the one of higher mammals..

Murine infection models and organoid culture technology contributed to better understand ZIKV neurotropism and neurovirulence. Infected adult mice or pups (immunodeficient or immunocompromised in most of the studies) show accumulation of ZIKV in both the brain and the spinal cord in addition to other organs such as liver, testes and spleen.^3,6,7,27^ However, the individual contribution of ZIKV proteins to neurovirulence was never addressed *in vivo* in vertebrate models. Furthermore, monitoring the physiology of NPCs in the whole brain of vertebrates, especially in transgenic mammalian models remains challenging because of high costs, invested time, ethical considerations, access to specific cell types inside the brain for imaging and omic analysis, and limited genetic plasticity associated to potential embryonic lethality. Thus, alternative models more conducive to the study of early development of the ZIKV-infected brain are required.

The zebrafish has emerged as a powerful and cost-effective tool for studying neurological diseases relevant to humans.^28,29^ The zebrafish embryo (1-to-2 day-old) and larva (3-to-6 day-old) are ideally suited to examine neurodevelopment in the brain as the neural population, connectivity and axon tracts as a whole are preserved during experimentation. The optically transparent fish represents an exquisite *in vivo* toolbox enabling easy imaging of the brain in transgenic animals, loss- or gain-of-function genetic approaches, behavioral tests to examine changes in motor activity, and the ease to use a theoretically unlimited number of animals. Furthermore, the larval zebrafish brain shares basic neuroanatomical layout with that of mammals.^30^ Moreover, zebrafish is permissive to several human viruses (*e.g.,* norovirus, herpes simplex virus, Rift Valley fever virus^31–33^) and phenocopies several human neuropathological diseases such as amyotrophic lateral sclerosis, fronto-temporal dementia and CHARGE syndrome.^28,29,34–36^

In this study, we have established a novel zebrafish-based ZIKV infection model to study viral neuropathogenesis *in vivo*. In this system, larvae infected with ZIKV exhibited severe morphological defects during their development. ZIKV infection was detected in brain NPCs, which correlated with a decrease in their abundance and in head size, as well as drastic mobility impairments and induction of apoptosis in the brain, thus phenocopying the disease in mammals. The transcriptomic analysis of NPCs isolated from whole larvae revealed that ZIKV downregulated the expression of the glutamate transporter *vglut1*, which was associated with an altered network of glutamatergic neurons in the brain. In contrast, ZIKV infection increased cell survival, apoptotic and differentiation regulation factors, such as *pim2*, *cbx7a*, and the components of the activator protein 1 (AP-1) family members (*jun, junB, fosab*). Importantly, the sole expression of ZIKV protein NS4A in the larvae fully recapitulated the morphological defects observed in infected animals. By mimicking several phenotypes and symptoms in humans caused by ZIKV, our innovative animal model provides a unique and unprecedented access to the ZIKV-infected brain of vertebrates to further investigate the host and viral determinants of ZIKV neuropathogenesis *in vivo*. Given the efficiency of known anti-ZIKV drugs in this system, it will also serve as a suitable platform for medium-throughput drug screening of antiviral molecules *in vivo*.

## RESULTS

### Zika virus replicates efficiently in zebrafish larvae and induces morphological defects

To establish a new *in vivo* model for ZIKV infection, we microinjected ∼45 infectious virus particles of the ZIKV H/PF/2013 strain in the yolk of zebrafish embryos within the two first hours following fertilization to be consistent with an infection during early pregnancy (Figure 1A). Embryos injected with DMEM (vehicle, mock) were used as reference controls. At 3 days post-fertilization (dpf), zebrafish larvae were assessed for changes in viability and morphology. We observed that 3 dpf ZIKV-infected larvae exhibited a marked decrease of approximatively 65% in the survival rate compared to controls (Figure 1B). ZIKV-infected larvae also showed drastic developmental defects at 3 dpf (Figures 1C and 1D). More specifically, ∼80% of the living larvae exhibited mild or severe morphological defects, ranging from curved and shorter tail, oedema to ovoid morphology. Importantly, a significant reduction of 20% in the head area was noted following infection with ZIKV compared to the controls (Figure 1E), which is reminiscent of newborn microcephaly in humans. Remarkably, injection of DENV, a non-neurovirulent flavivirus did not induce any apparent morphological defects particles (except heart oedema in some larvae). Moreover, in contrast to ZIKV, no differences in viability or morphology were observed between mock- and DENV-infected larvae (Figures 1B-1E). Altogether, these data suggest that the defects induced by ZIKV injection in zebrafish larvae are specific to this virus.

**Figure 1.**
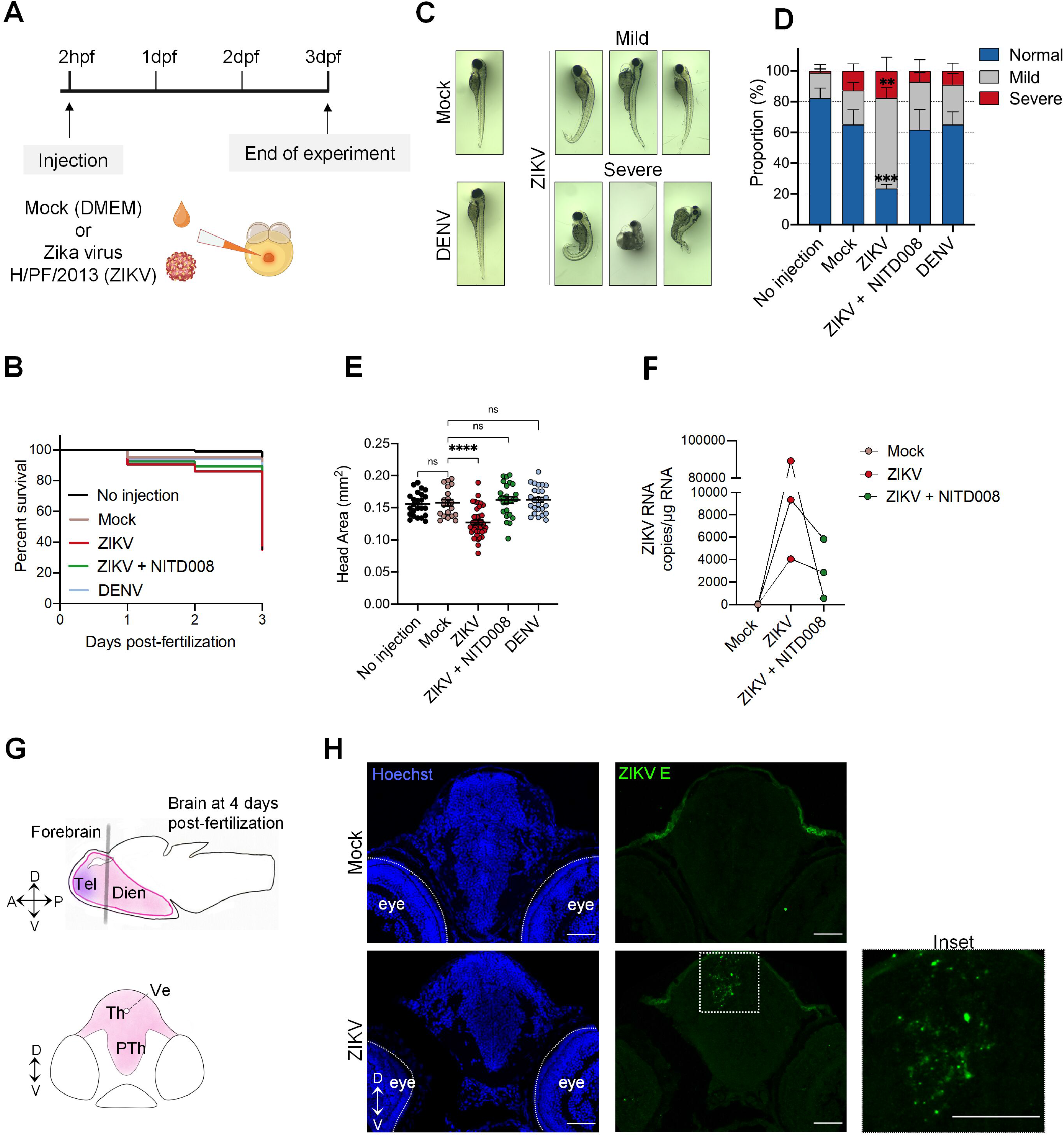
Zika virus replicates efficiently in the zebrafish model and induces morphological defects. (A) Schematic experimental design. Cell medium (mock), DENV viral particles (serotype 2, strain 16681s) or ZIKV viral particles (strain H/PF/2013) were microinjected in the zebrafish yolk at 2 hours post-fertilization (hpf). (B) Survival curve over 3 days post-fertilization (dpf) of mock-infected (*n*=84), ZIKV-infected (*n*=104), ZIKV-infected and treated with NITD008 (*n*=94) and DENV-infected (*n*=89) larvae (*N*=3). (C) Representative pictures of microinjected larvae at 3 dpf. ZIKV infection induced both mild and severe developmental phenotypes. (D) Quantification of the proportion of larvae with the different phenotypes (No injection, *n*=64; Mock, *n*=54; ZIKV, *n*=67; ZIKV+NITD008, *n*=62; DENV, *n*=55. *N*=3). Data are shown as means ± SEM. *** P ≤ 0.001; ** P ≤ 0.01; 2-way ANOVA. (E) Head size at 3dpf of the larvae from (D). Data are shown as means ± SEM. **** P ≤ 0.0001; ns = not significant; one-way ANOVA. (F) ZIKV RNA levels in whole larvae at 3 dpf were determined using ddPCR. (G) Schematic representation of a zebrafish brain at 4 days post-fertilization. The forebrain is shown. Gray line represents the localization of the transverse sections. Th and PTh are areas of the diencephalon, a division of the forebrain. Dien= diencephalon; Tel= telencephalon; Th= thalamus; PTh= prethalamus; Ve= venticle. A= anterior; P= posterior; D= dorsal; V= ventral (H) Transverse brain cryo-sections of 4 days post fertilization mock-injected or ZIKV-injected larvae are shown. Sections were stained with anti-ZIKV E antibody (green) and analyzed by confocal microscopy. Nuclei were labeled with Hoechst. Scale bars = 50 μm. *n* represents the number of fish; *N* represents the number of independent experimental repeats.

Toward demonstrating that ZIKV efficiently replicates in zebrafish, NITD008, a nucleoside analog which inhibits ZIKV NS5 RNA polymerase and thus, viral replication^37^, was added to the fish water at 4 hours post-infection. Total RNA was extracted from whole larvae and viral RNA levels were measured using droplet digital PCR (ddPCR). Viral RNA was readily detected at 3 dpf but its levels were reduced up to 150-fold when larvae were treated with NITD008, demonstrating that the virus replicates in the animal (Figure 1F). Most importantly, NITD008-treated infected larvae had similar survival rate and morphology as uninfected fish (Figures 1B-1E), suggesting that the observed defects were solely due to ZIKV replication, and not indirectly caused by innate immunity-dependent inflammatory responses. Of note, RT-qPCR with total RNA extracted from entire infected larvae failed to detect any induction of interferon (*ifnΦ1*) and *rig-I* mRNAs, which is an interferon-stimulated gene (Figure S1A). These data along with the fact that DENV injection did not induce any major defects, strongly indicate that morphological defects are specific to ZIKV replication.

To test whether ZIKV is neurotropic in zebrafish, larvae brain cryosections were immunolabelled at 4 dpf with a panflaviviral anti-envelope (E) antibody. Infection foci were specifically detected in the developing forebrain and midbrain of ZIKV-infected animals compared to control animal with high signal intensity in the thalamus of the diencephalon, an area surrounding the brain ventricle in the forebrain (Figures 1G and 1H). This indicates that ZIKV is neurotropic in the larva even if the injection was performed at a time at which cells were pluripotent.^38^

Taken together, our findings unambiguously demonstrate that zebrafish larva is permissive to ZIKV which replicates in the developing brain, inducing phenotypes highly reminiscent of those observed in infected human fetuses. This strongly suggests that the observed ZIKV-induced morphological and head defects are caused by ZIKV brain neurovirulence as in mammals.

### ZIKV infection causes mobility defects

While it is clear that ZIKV targets the CNS as in human, we have further investigated to which extent the zebrafish model mirrors the human disease in terms of neurovirulence and severity of the symptoms. Previous epidemiological studies showed that motor impairment is associated to CZS in human.^39,40^ Of note, locomotor activities in zebrafish are closely linked to brain function integrity, to visual development, to muscle activity, and more importantly to nervous system development. ^41–44^ Therefore, we hypothesized that motor activity can be used as a readout of nervous system development and brain abnormalities in zebrafish. To challenge this hypothesis, we performed a touch-elicited escape behavioral assay using live animal imaging at 2 and 3 dpf in ZIKV-infected fish and controls. Most ZIKV-injected animals with mild morphological defects were unable to flee at 2 dpf, displaying abnormal circular swimming patterns upon stimulation when compared to controls (Video S1). At 3 dpf, very little movement, if any was observed for the infected larvae (Video S2). Very strikingly, NITD008 treatment seemingly reverted these swimming defects (Videos S1-2). This observation is consistent with the inhibition of ZIKV replication following treatment with this drug (Figure 1F).

To rigorously assess the impact of ZIKV infection on the motor system, we used the Daniovision, an automated observation chamber which allows the quantitative analysis of motor behaviors (Figure 2A). At 4 days post-fertilization, ZIKV-infected larvae exposed aberrant swimming behavior. Indeed, zebrafish infected larvae had little to no movement (Figure 2B). Using this technique, we showed that the distance swum following infection was severely reduced by ∼25-fold when compared to control larvae (Figure 2C), consistent with an increase in immobility time (Figures S1B and S1C). In accordance with live imaging observations, NITD008 partially restored the swimming capacity (Figures 2B and 2C). Moreover, DENV injection did not induce any mobility defects (Figures S1B and S1C), suggesting that the phenotypes are specific to ZIKV replication.

**Figure 2.**
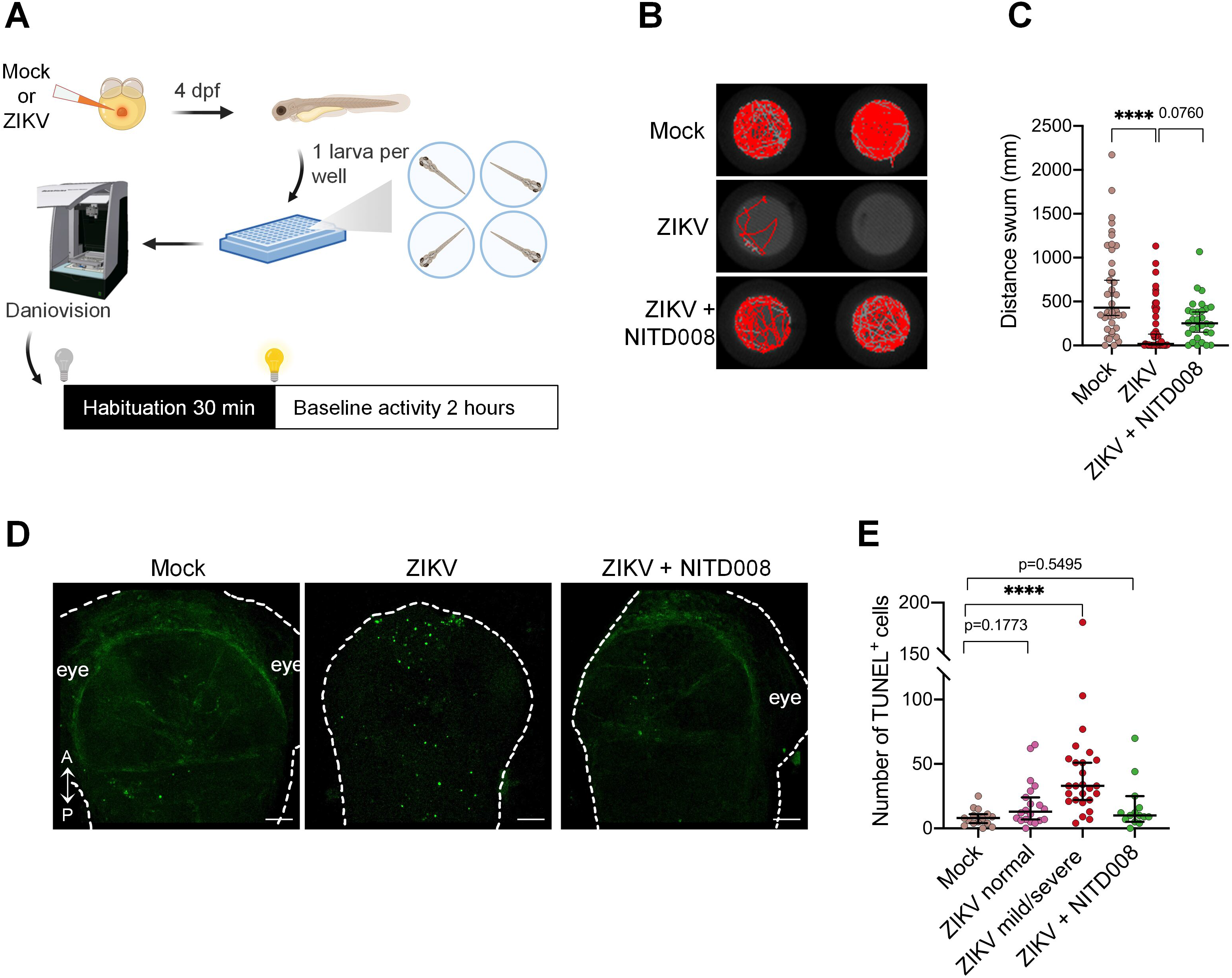
Locomotor defects and brain cell death in zebrafish larvae following ZIKV infection. (A) Schematic experimental setup and behavioral analysis. (B) Representative swimming tracks of control (mock), ZIKV-infected (ZIKV), and ZIKV-infected and NITD008-treated (ZIKV+NITD008) larvae at 4 days post-fertilization. (C) The distance moved by the larvae was assessed using the DanioVision device (mock, *n*=40; ZIKV, *n*=45; ZIKV+NITD008, *n*=30. *N*=3). Data are shown as median ± 95 CI. **** P ≤ 0.0001; Kruskal-Wallis test. (D) TUNEL staining at 2 days post-fertilization showing cell death in the developing brain following injection in zebrafish embryo. A = anterior; P = posterior. Scale bars = 50 μm (E) The number of TUNEL+ cells was quantified (Mock, *n*=15, *N*=3; ZIKV normal, *n*=20, *N*=3; ZIKV mild/severe, *n*=26, *N*=3; ZIKV+NITD008, *n*=14, *N*=2). Data are shown as median ± 95 CI. **** P ≤ 0.0001; Kruskal-Wallis test. *n* indicates the number of fish; *N* represents the number of independent experimental repeats.

### ZIKV induces cell death in the developing brain of zebrafish

Earlier studies have shown that ZIKV infection leads to cellular death *in vivo* and *in vitro*.^5,6^ Considering ZIKV neurotropism, the reduction in head area and the locomotor defects in our zebrafish larva model, we sought to investigate further the extend of neurovirulence by evaluating ZIKV-induced apoptosis in the developing brain. At 2 days post-fertilization, whole fixed larvae were subjected to terminal deoxynucleotidyl transferase-mediated dUTP nick end-labeling (TUNEL) assays to detect apoptotic cells by confocal microscopy. Subsequent quantification of the TUNEL-positive cells revealed an overall 3-fold increase in the number of apoptotic cells in the developing brain following viral infection, as compared to the control (Figures 2D and 2E). More precisely, ZIKV-injected larvae displaying mild or severe morphology defects exhibited a ∼4-fold increase in brain cell death compared to the control (Figure 2E). Notably, even ZIKV-injected larvae with seemingly normal morphology displayed an increased number of apoptotic cells when compared to the control group, although the difference was not statistically significant (Kruskal-Wallis test, p-value=0.1773). Interestingly, ZIKV-injected larvae treated with NITD008 had similar cell death level than the control (Figures 2D and 2E). These results demonstrate that ZIKV replication induces cell death in the developing brain.

### ZIKV infects neural progenitor cells and induces their depletion in zebrafish developing brain

Among other cell types, NPCs were described to be a major target of ZIKV in newborns. Thus, we investigated the impact of ZIKV infection on NPC abundance and their distribution in the brain. First, we took advantage of the transgenic line Tg(*gfap*:GFP), which allows visualization and quantification of NPCs at 1 day post-fertilization. In this transgenic line, the native (*i.e.* non-fused) green fluorescent protein (GFP) is expressed under the transcriptional control of the glial fibrillary acidic protein (*gfap*) promotor, a marker of NPC at this early time point of brain development.^45,46^ First, we confirmed that ZIKV injection in these transgenic embryos recapitulates the morphological defects observed in wildtype larvae (Figure S2A). Next, Tg(*gfap*:GFP) embryos were infected with ZIKV or left uninfected and 24 hours post-fertilization, *i.e.*, shortly after neurogenesis induction^47^, whole embryos were dissociated into single cells. GFP-positive cells were quantified by flow cytometry in the presence of fluorescent beads, allowing to normalize for dissociation efficiency, and unbiased cell counting as described before (Figure 3A).^48^ Reverse transcription PCR (RT-qPCR) on sorted GFP+ cells confirmed high endogenous expression of *nestin* and *gfap* mRNAs, hallmarks of NPCs at 1 dpf, when compared to GFP^-^ cells (Figure S2D), confirming selective expression of GFP in the neural progenitor cells.^49,50^ Strikingly, ZIKV infection induced a decrease in the abundance of GFP^+^ cells, *i.e.*, NPCs, compared to the control (Figure 3B). Particularly, we observed a mean 64.4%-fold decrease of NPC (one-way ANOVA; p ≤ 0.01) when the phenotype was mild or severe (Figure 3B). We additionally investigated an additional transgenic line Tg(*nestin*:GFP), allowing the detection of neuronal progenitors, a subclass of NPCs. Similar to wildtype line larvae, ZIKV injection induced morphological defect in this line (Figure S2B). Consistently with the results obtained with Tg(*gfap*:GFP), the number of neuronal progenitor cells (GFP^+^) was markedly decreased in Tg(*nestin*:GFP) (Figure S3C). These data unambiguously demonstrate a loss of NPCs in the whole embryo following ZIKV-infection.

**Figure 3.**
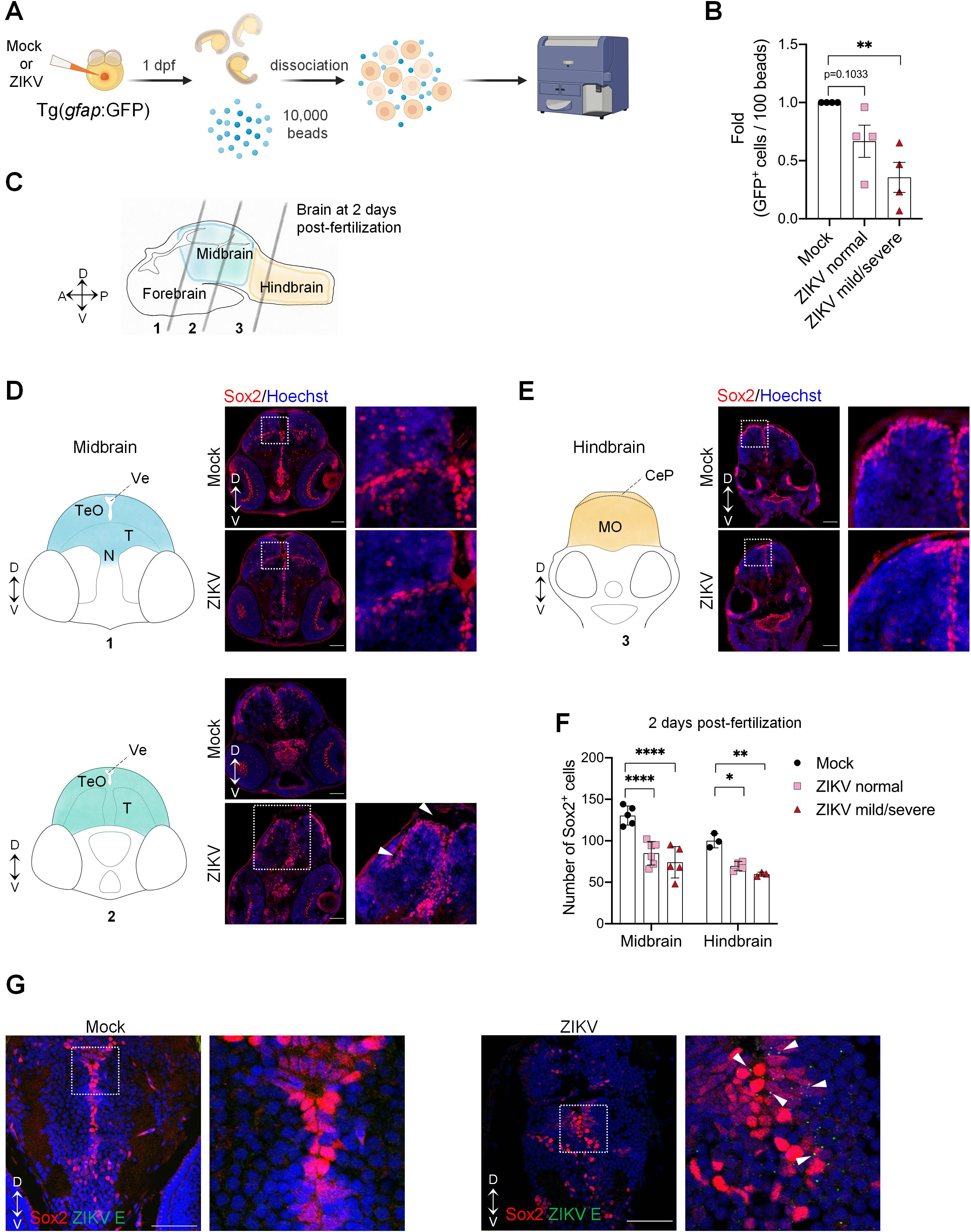
Zika virus targets neural progenitor cells and induces neuropathogenesis in zebrafish larvae. (A) Schematic experimental design. At 1 day post-fertilization, ZIKV infected or uninfected whole transgenic Tg(*gfap*:GFP) embryos were dissociated in the presence of 10,000 fluorescent normalizing beads. Single cells and beads were counted by flow cytrometry. (B) Number of GFP^+^ cells counted per 100 beads (*N*=4). Data are shown as means ± SEM. ** P ≤ 0.01; one-way ANOVA. (C) Schematic representation of a zebrafish developing brain at 2 days post-fertilization showing the three areas of the brain: forebrain, midbrain, and forebrain. Gray lines represent the localization of the transverse sections. D= dorsal; V= ventral; A= anterior; P= posterior. (D-F) Number of neural progenitor cells (Sox2^+^ cells) in the midbrain (D) and the hindbrain (E) of 2 days post-fertilization mock-injected or ZIKV-injected embryos. TeO, N, and T are areas of the midbrain while MO is an area of the hindbrain. TeO= tectum opticum; T= midbrain tegmentum; N= region of the nucleus of medial longitudinal fascicle; Ve= ventricle; MO= medulla oblongata; CeP: Cerebellar plate. D= dorsal; V= ventral. Data are shown as means ± SEM. **** P ≤ 0.0001; ** P ≤ 0.01; * P ≤ 0.05; Two-way ANOVA. Scale bars = 50 μm. (G) Confocal microscopy of brain section from mock-injected and ZIKV-injected embryos at 4 dpf. Cells were co-immunostained with anti-Sox2 and anti-ZIKV E. Cell nuclei were counterstainted with Hoechst. Scale bars = 50 μm.

We aimed to gain more insight in NPC distribution and abundance in different brain regions following ZIKV infection. Brain cryosections of ZIKV-infected fish and controls at 2 and 4 dpf were immunolabeled for Sox2, another marker for neural progenitor cells (Figures 3C-3F and S3). ^47^ In agreement with earlier reports, multiple layers of Sox2*^+^* cells could be identified in the tectum opticum of control animals (Figure 3D) at 2 dpf, while ZIKV-infected embryos displayed a significantly thinner layer (Figure 3D top inserts).^51^ At this time, Sox2^+^ cells were also located in periventricular zones, which are in direct contact with the midbrain and hindbrain ventricles. More precisely, they were lining the walls of the midbrain midline ventricle (Figure 3D bottom) and were in the ventricular surface of the cerebellar plate, and in medulla oblongata (Figure 3E). In control brain, NPCs are distributed along the ventricular surfaces, following a dorsomedial-dorsolateral distribution as previously described.^51^ Compared to mock, Sox2^+^ cells in ZIKV-infected embryo were distributed to a lesser extent on the lateral sides of the midbrain and hindbrain, suggesting a defect in positioning (Figures 3D and 3E). Quantification of Sox2^+^ cells in the midbrain at 2 dpf revealed a significant decrease in the number of NPCs of ZIKV-infected fish displaying both normal, and mild/severe phenotypes (Figure 3F). Consistently, depletion of NPCs was also observed in the midbrain at 4 dpf (Figures S3A-S3B and S3D), and in the hindbrain at 2 and 4 dpf (Figures 3E-3F and S3C-S3D). This demonstrates that ZIKV infection specifically targets the pools of NPCs, reducing their number and density in different parts of the developing brain and interfering with their positioning. Of note, brains from ZIKV-infected larvae brain displayed a dilated ventricle compared to control (Figure 3D, bottom, white arrows), resembling the ventriculomegaly and hydrocephaly observed in ZIKV-infected human newborns.^52,53^

In order to confirm that impairment of NPCs resulted from ZIKV infection, 4 dpf brain sections were co-stained with antibodies directed against Sox2 and ZIKV E. Infected NPCs mostly accumulated in periventricular regions rich in Sox2*^+^* cells (Figure 3G). Precisely, almost all infected cells were positive for Sox2, demonstrating that ZIKV primarily infects NPCs. Altogether, these observations demonstrate that ZIKV infection of zebrafish at early stages of development closely resembles the pathophysiology of human infections upon vertical transmission with respect to the physiology, neurodevelopmental sequalae and specific perturbation of the NPCs pool in terms of abundance and distribution.

### ZIKV infection modulates the expression of genes involved in cellular proliferation, differentiation, and apoptosis in NPCs

To identify the specific molecular mechanisms and viral targets underlying ZIKV-induced impairment of NPC abundance, we investigated the impact of ZIKV infection on the transcriptional landscape of NPCs during development *in vivo*. NPCs were isolated from ZIKV-infected and control Tg(*nestin*:GFP) embryos at 1 dpf using fluorescence-activated cell sorting (FACS). As in wildtype line larvae, ZIKV induced morphological defects in Tg(*nestin*:GFP) (Figure S2B). We confirmed that NPC-specific endogenous *nestin* and *gfap* mRNAs were enriched in isolated GFP^+^ cells, compared to GFP^-^ cells (Figure S2E). RNA sequencing of isolated NPCs revealed significant transcriptional modulation of 199 genes in ZIKV-infected NPCs, with 104 and 95 genes significantly up- and down-regulated, respectively (Table S1; P<0.01; Log_2_FoldChange > 0.7 or < −0.7).

Interestingly, mRNAs encoding for cell proliferation and apoptotic regulator factors such as *pim2* and AP-1 transcription factor members *junba, jun and fosab* were increased. *Chromobox homolog 7a (cbx7a)* expression, which inhibits differentiation, axon growth and axon regeneration was also increased by 6.7-fold (p-value= 4.59 x 10^-7^, Figure 4A).^54,55^ These results suggest a ZIKV-induced rewiring of zebrafish NPC survival and neuronal maturation networks, consistent with those reported for mammalian NPCs.^11^

**Figure 4.**
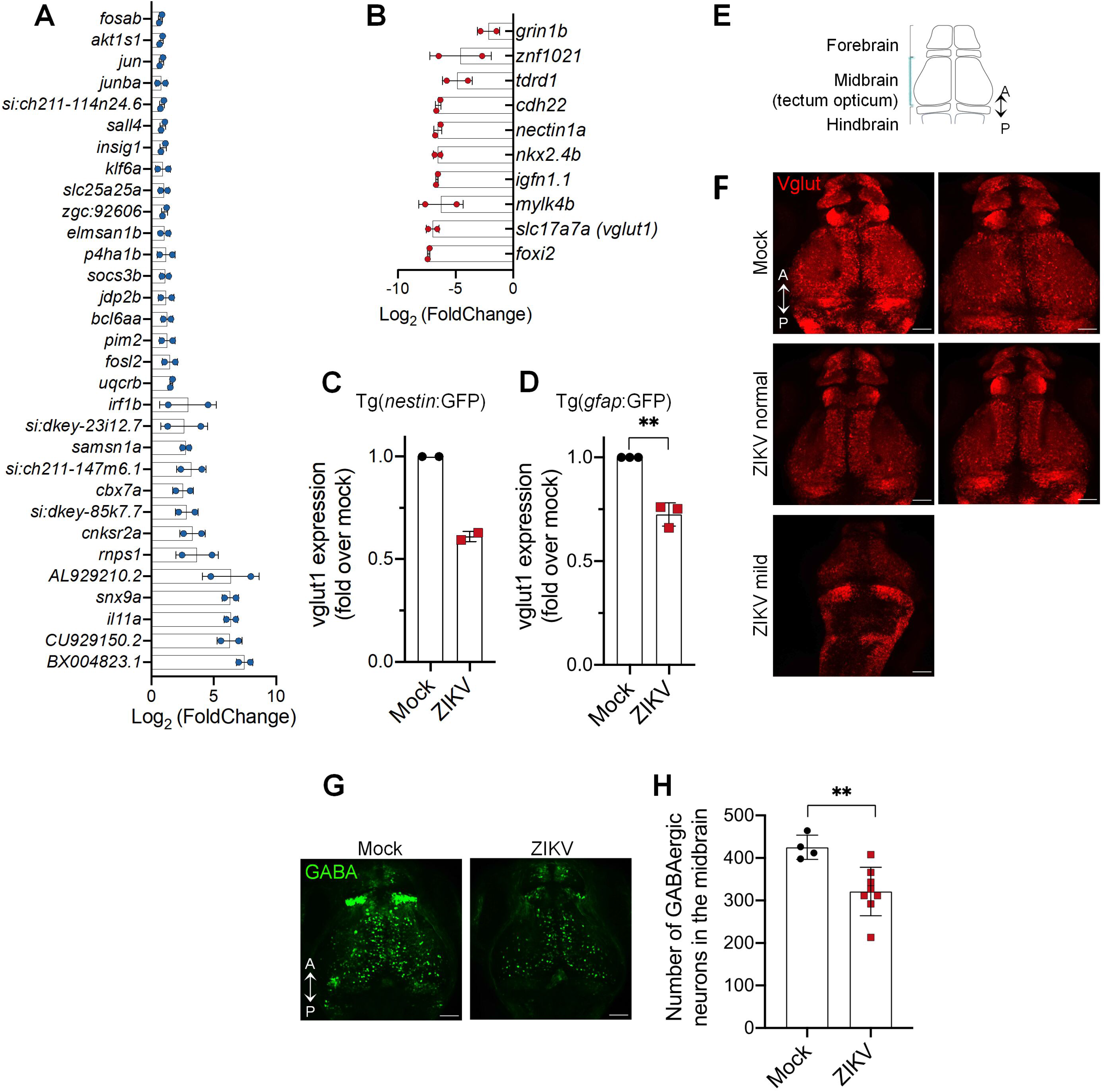
Impact of ZIKV infection on zebrafish larva transcriptome. (A and B) Tg(*nestin*:GFP) embryos were infected with ZIKV or left uninfected. At 1 dpf, NPCs (*i.e*., GFP+ cells) were isolated from whole larvae using FACS. NPC transcriptome was analyzed using RNA sequencing. mRNA expression levels were compared to that of NPCs isolated from uninfected control larvae. The most significantly upregulated (A) and downregulated (B) genes are shown (*p* < 0.01) (Mock, *n*=30; ZIKV, *n*=31. *N*=2). (C and D) *vglut1* gene expression in NPCs isolated from infected larvae compared to uninfected control larvae using two transgenic cell lines Tg(*nestin*:GFP) (C) and Tg(*gfap*:GFP) (D) was analyzed by ddPCR. (E) Schematic representation of 3 dpf larval brain from dorsal view. A = anterior; P = posterior. (F) Confocal microscopy from whole brain 3dpf larvae showing the glutamatergic neurons network. Scale bars = 50 μm. (G-H) Confocal microscopy from whole brain 3dpf larvae showing the glutamatergic neurons network. (H) Quantification of (G). Data are shown as means ± SD. ** P ≤ 0.01; Student’s t-test. Scale bars = 50 μm. *n* indicates the number of fish; *N* represents the number of independent experimental repeats.

Most notably, one of the most affected genes was *vglut1*, displaying a 130-fold reduced expression upon ZIKV infection (p-value= 2.39 x 10 ^-5^, Figure 4B). ddPCR analysis of NPCs isolated from infected Tg(*nestin*:GFP) and Tg(*gfap*:GFP) embryos showed a decreased expression of *vglut1*, thus confirming the expression profile of *vglut1* in the RNA sequencing results (Figures 4C and 4D). *Vglut1* mRNA encodes the glutamate transporter, which is required for the release of this neurotransmitter by presynaptic excitatory neurons. It is thus a *bona fide* marker of glutamatergic neurons. Additionally, the abundance of *glutamate receptor ionotropic NMDA 1b* (*grin1b*) mRNA was also decreased (Figure 4B). The protein encoded by this gene is a subunit of the NMDA glutamate receptor. *GRIN1* mutations, and more broadly defects in the functionality of these glutamatergic neurons, are typically associated with cognitive impairments, epilepsy, microcephaly, muscular tone abnormalities, and behavior issues.^56–59^ Interestingly, such impairments are a sequelae of children born with the congenital Zika syndrome.^60,61^ To test whether *vglut1* and *grin1b* mRNA levels decrease correlates with a glutamatergic network alteration, we characterized this neuron population using the Tg(*dlx5a/6a*:GFP;*vglut2*:RFP) fish line, in which glutamatergic neurons express red fluorescent protein (RFP) while GFP is produced in GABAergic neurons. Whole brain imaging identified reduced RFP intensity signal in all parts of the brain *i.e*. forebrain, midbrain and hindbrain following ZIKV infection (Figures 4E and 4F) demonstrating that ZIKV disrupts the excitatory glutamatergic network.

In addition to cognitive impairments, epilepsy/seizure are a frequent occurrence in children with CZS.^62,63^ Recently, alteration of GABAergic interneurons were shown to be associated with hyperactivity in zebrafish, γ-Aminobutyric acid GABA being the primary inhibitory neurotransmitter in the CNS.^35^ The abundance of GABAergic neurons (GFP^+^ cells) was significantly reduced in Tg(*dlx5a/6a*:GFP;*vglut2*:RFP) larvae following ZIKV infection compared to control, although it is noteworthy that no change in the expression of *dlx* genes (*dlx1a, dlx2a, dlx5a*, and *dlx6a*) involved in the specification of GABAergic neurons was noted (Figures 4G, 4H and Table S1). Collectively, our results suggest that the cognitive impairment and neurodevelopmental disorder induced by ZIKV are partly due to loss of glutamatergic and GABAergic neurons likely caused by an alteration of neurogenesis.

### NS4A is a major viral determinant of ZIKV neuropathogenesis

The viral determinants of ZIKV neurovirulence remain poorly understood. A previous study has shown that ubiquitous expression of ZIKV NS4A protein in drosophila induces severe neurodevelopmental defects including a reduced larval brain volume and apoptosis in neurons.^2526^ However, drosophila is an invertebrate model whose brain development substantially differs from that of humans. Therefore, we assessed neurovirulence potential of NS4A in the zebrafish larva, a vertebrate model closely resembling brain development in higher vertebrates and humans. We also included in the analysis NS4B since it interacts with NS4A and was shown to inhibit neurogenesis *in vitro*.^11^ To confirm that *in vitro* transcribed RNAs can be correctly translated by the zebrafish cellular machinery, we microinjected a full-length *in vitro* transcribed ZIKV RNA genome (vRNA). vRNA injection fully recapitulated the morphological defects observed in infection (Figures S4A-S4D), demonstrating that this approach is in principle appropriate to investigate the impact of individual viral proteins expression. *In vitro* transcribed RNA encoding NS4A, NS4B (with the 2K signal peptide), or NS4A-(2K)-NS4B (NS4A precursor) were microinjected into 1 cell stage zebrafish embryos (Figure 5A). We then examined the effects of NS4A, NS4B, and NS4A-NS4B precursor expression on the general morphology. Embryos were analyzed for morphology and motor activity. Strikingly, like for ZIKV-infected larvae, NS4A-injected animals displayed severe mobility restrictions (Video S3). NS4A expression caused morphological defects at 3 dpf such as curved or shortened tails as compared to the water injection control. Indeed, 60% of those embryos had mild or severe defects (Figures 5B and 5C). More importantly, NS4A expression led to 20% decrease of the head area (Figure 5D). This correlated with an increase of apoptosis in the brain at 2 dpf as measured using TUNEL assays (Figure 5F and 5G). In contrast, NS4B or NS4A-NS4B expression did not lead to an increase in morphological defects compared to the controls (Figures 5B-5D), demonstrating that the phenotypes are specific to mature ZIKV NS4A. It is noteworthy that hatching of embryos injected with NS4A was seriously affected, with 40% of NS4A-injected larvae in their chorion at 3 dpf, a time point by which hatching has normally already occurred (Figure 5E).^64^ Taken together, our results unequivocally demonstrate that NS4A is sufficient to trigger morphological and locomotor defects closely resembling those observed in infection and to induce neurovirulence, constituting an attractive drug target for therapeutic development.

**Figure 5.**
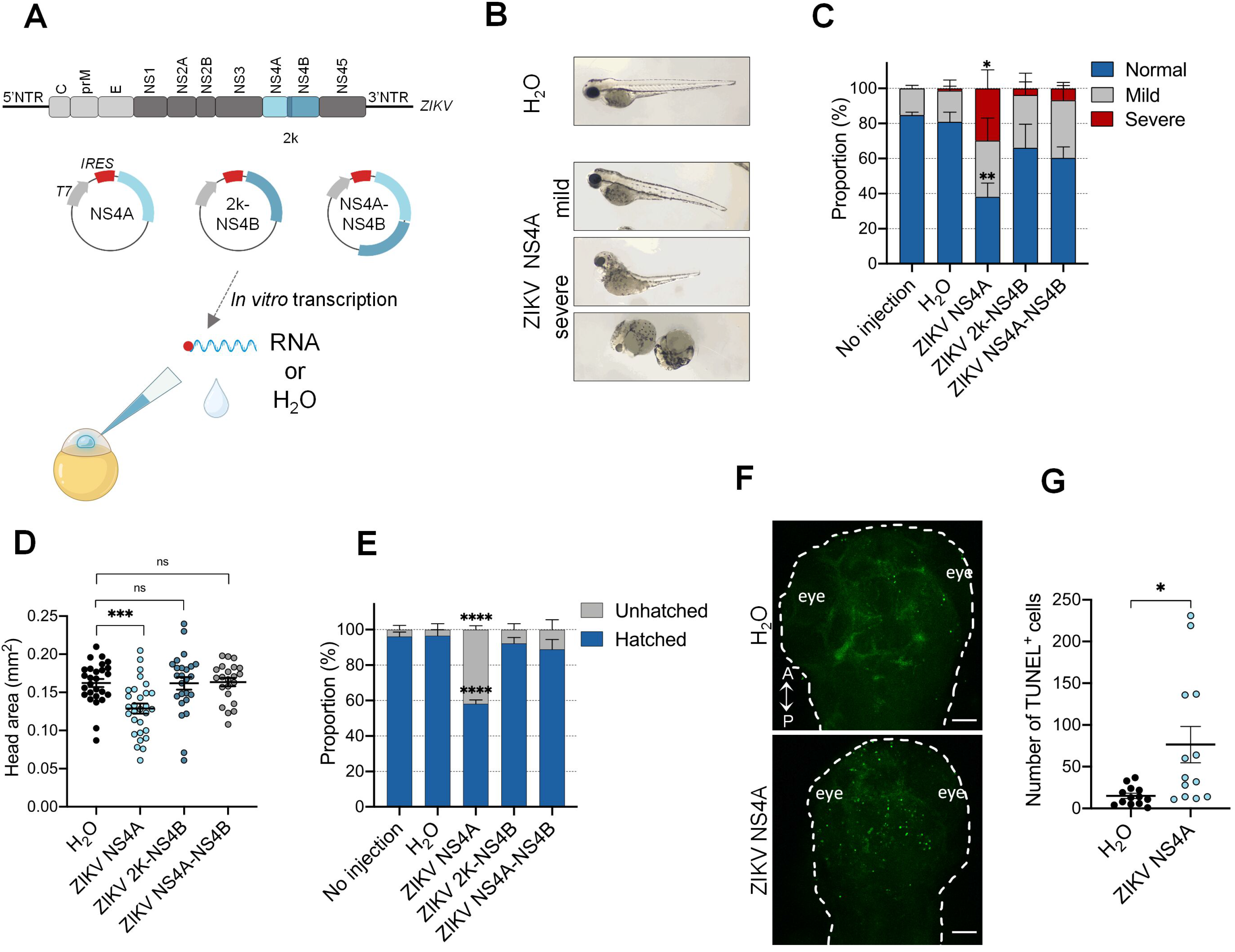
ZIKV NS4A is a major viral determinant in ZIKV pathogenesis *in vivo*. (A) Schematic experimental design. ZIKV genome is represented. *In vitro* transcription was performed on plasmid coding for NS4A, 2K-NS4B, and NS4A-NS4B. NTR = non-translated region. IRES = internal ribosomal entry site. *In vitro* transcribed NS4A, 2K-NS4B, NS4A-NS4B RNAs were injected at one-cell stage at 1 hour post-fertilization. H_2_O was used as reference control. (B) Representative pictures of microinjected larvae at 3 days post-fertilization. NS4A expression induced both mild and severe developmental phenotypes. (C) Proportion of larvae with different phenotypes at 3 days post-fertilization (No injection, *n*=68; H_2_O, *n*=53. ZIKV NS4A, *n*=62. ZIKV 2K-NS4B, *n*=62. ZIKV NS4A-NS4B, *n*=63. N=3. Data are shown as means ± SEM. **** P ≤ 0.0001; ** P ≤ 0.01; * P ≤ 0.05; ns: non-significant; two-way ANOVA. *N*=3. (D) Head area quantification of mock (*n*=27), ZIKV NS4A-(*n*=29), ZIKV 2K-NS4B-(*n*=25) and NS4A-NS4B-(*n*=22) injected larvae at 3 days post-fertilization. Data are shown as means ± SEM. *** P ≤ 0.001; ns: non-significant; one-way ANOVA. *N*=2. (E) Hatching rate at 3 days post-fertilization of larvae analyzed in (C). (F and G) TUNEL staining at 2 days post-fertilization (F) and cell death quantification (G) in the developing brain following ZIKV NS4A injection (*n*=13) compared to H_2_O-injected (*n*=13) zebrafish embryo. *N*=2. A= anterior; P= posterior. Scale bars = 50 μm. *n* indicates the number of fish; *N* represents the number of experimental repeats.

## DISCUSSION

Despite multiple investigations using existing immunocompromised murine models and recently developed organoid culture models, the viral and host determinants driving ZIKV neuropathogenesis are still poorly understood. Our study unveils the zebrafish larva as a vertebrate model for ZIKV infection with neurological phenotypes comparable to those in humans. Taking advantage of transgenic lines with specifically labelled neural cell populations combined with molecular and cellular analysis, we uncovered several host and viral determinants of ZIKV pathogenesis.

Upon injection in the yolk at 2 hours post-fertilization, ZIKV infects and replicates efficiently in the zebrafish developing brain, notably in NPCs, a known target of ZIKV in humans. Despite cells being pluripotent at the time of injection, ZIKV is primarily detected in neural progenitor cells at 4 days post-fertilization, indicating a specific cellular tropism. However, ZIVK infection was not restricted to neural progenitor cells since we detected a fraction of ZIKV envelope in Sox2-negative cells. These cells could be astrocytes, oligodendrocytes or microglial cells, which are also permissive to ZIKV ^65^, or infected NPCs that have initiated their differentiation into post-mitotic neurons and thus, lost *sox2* expression. Staining with specific markers of other neural population markers such as Neurod1, a marker for intermediate neurons or Elavl3, a marker of immature neurons, may be used in future studies to better discriminate ZIKV-infected cell populations. Future studies involving single-cell sequencing will allow a better understanding of ZIKV cellular tropism and neuronal maturation, especially considering that previous studies reported that ZIKV impairs neuronal differentiation, consistent with our RNA sequencing data.

Injection of ZIKV at very early stage causes developmental defects in the zebrafish larvae characterized by morphological defects, mobility impairments, and more importantly, by a decrease in head area. Infection with DENV did not lead to any obvious abnormalities in the general development, except discrete heart oedema in some scarce occurrences. This strongly suggests that these phenotypes are specific to ZIKV infection and are not due to inflammation or antiviral host responses. Treatment of ZIKV-infected animals with the NS5 polymerase inhibitor NITD008 systematically reverted the phenotypes. It unambiguously demonstrates that productive ZIKV replication is required to induce developmental defects. Such experiments additionally represent a proof-of-principle that zebrafish can be used for drug *in vivo* efficacy studies, enabling convenient and rapid testing of antiviral drug candidates before challenge in more elaborated, expensive, and time-consuming challenges in murine models. Moreover, considering that the drug was added to the water and efficiently taken-up by the larva, this model is ideal for *in vivo* medium throughput drug screening campaigns to identify novel antivirals, which would rely on ZIKV-induced defect reversion upon treatment. Our infection model, combined with behavioral assays using the Daniovision recording chamber, enables quantitative and automated drug testing, highlighting the scalability of our approach.

ZIKV replication led to increased cell death in the developing brain of the larva which is consistent with previous reports.^6^ Together with neural progenitor cells depletion, these could explain the decrease in head area observed in infected larvae. It is noteworthy to mention that increased apoptosis in the brain and decrease NPC abundance were also observed in some infected animal without general physical defaults (Figures 2E, 3B, 3F and S3D, “normal” animals). In human, non-microcephalic infants with intrauterine exposure to ZIKV can exhibit neurodevelopmental delays such as cognitive and language impairments. ^66,67^ More studies of those infected fish could allow a better understanding of sequalae and long-term consequences of ZIKV infection. Our transcriptomic analysis of NPCs from infected larvae reveals that one of the most affected genes was *vglut1* with a decrease in mRNA levels of ∼130-fold. *vglut1* mRNA encodes the vesicular glutamate transporter type 1 and is a specific biochemical marker of glutamatergic neurons and glutamatergic synapses. Several evidence suggest an inhibitory/excitatory imbalance and particularly abnormalities in excitatory glutamatergic neurons and synapses in neurodevelopmental disorders including intellectual disabilities and epilepsy, which are sequelae of children born with the congenital Zika syndrome.^68–72^ This imbalance correlates with severe mobility defects in both infected zebrafish and affected children. Interestingly, no modulation of genes involved in immune antiviral responses could be observed, suggesting that ZIKV-induced pathological sequelae are not driven by interferon signaling. It is thus plausible that the neurodevelopmental defects induced by ZIKV are partly due to a perturbation of the development, and functionality of the mature glutamatergic and/or GABAergic neuronal network. This may pave the way to novel therapeutic approaches aiming at dampening the symptoms in patients by chemically targeting one of these two mature neuron types, depending on the sequalae.

Here, we demonstrate that ZIKV NS4A is a major determinant of ZIKV neurovirulence in vertebrates as its expression alone at the early stages of development closely recapitulates the phenotypes observed upon infection. Although previous studies in drosophila have shown that ZIKV NS4A expression reduces the size of the larval brain ^25,73^, this is the first time to our knowledge that it is shown *in vivo* in a vertebrate model whose neuroanatomy and neurodevelopment resemble that of humans. As in infection, NS4A expression induced morphological defects correlated with mobility impairment, a reduction in head size and the induction of apoptosis in the larval brain. In drosophila, NS4A induces microcephaly by inhibiting the host protein Ankle2 and its pathway. Moreover, a recent study reported that Ankle2 knock-out in zebrafish resulted in microcephaly and a decrease in the number of radial glial progenitor cells, suggesting a role in neurogenesis in this model.^74^ Since NS4A also associates with Ankle2 human orthologue, it will be interesting to evaluate whether this interaction regulates NS4A neurovirulence in zebrafish larva. In human cells, flaviviral NS4A is strictly required for vRNA replication and accumulates in viral replication organelles which host the vRNA replication complexes.^16,75–78^ Both mutagenesis and pharmacological approaches recently demonstrated that NS4A regulates the biogenesis of these replication factories.^12,17^ Most notably, Riva and colleagues have shown that ZIKV NS4A function in replication organelle morphogenesis can be specifically inhibited by the compound SBI-0090799.^17^ Moreover, NS4A inhibits the Akt-mTOR pathway and the growth of neurospheres.^11^ Finally, in the case of DENV, NS4A can dampen RIG-I-dependent interferon induction in cell culture^79,80^ suggesting that it contributes to neuropathogenesis by interfering with innate immunity *in vivo* although this remains to be addressed. Overall, these multiple roles of NS4A highlight that this viral protein constitutes an attractive target for the development of direct-acting agents. Such drug would provide a dual therapeutic benefit by inhibiting both its functions in viral replication and pathogenesis.

While it is clear that NS4A plays a critical role in ZIKV-induced neurodevelopmental defects, one cannot exclude the possibility of other viral determinants synergizing NS4A activity to promote ZIKV neuropathogenesis. NS4A and NS4B cooperate and suppress the Akt-mTOR pathway which leads to inhibition of NPC growth *in vitro* and stimulation of autophagy.^11^ However, in our system, the overexpression of NS4B or NS4A-NS4B precursor did not induce any notable phenotypes. Considering that in the case of DENV, NS4A and NS4B interact together^16^, NS4B might potentialize NS4A activity when co-expressed in zebrafish. Interestingly, one defect observed in ZIKV-injected larvae was body curvature, a typical phenotype of ciliopathy in zebrafish.^81,82^ It is thus tempting to speculate that this phenotype is likely due to ZIKV NS5 protein, known for inducing ciliopathy, forcing premature neurogenesis in chicken embryo and affecting motile cilia located in the brain in human fetal microcephalic tissue.^83^ This suggests that this viral protein also contributes to viral neuropathogenesis.^83^

In summary, our data unveil the zebrafish larva as a new versatile model of ZIKV pathogenesis since it recapitulates ZIKV neurotropism and CNS developmental defects observed in humans. ZIKV induces apoptosis and NPC depletion, which contributes to the development of microcephaly. Our study also provides important insights into host and ZIKV determinants of neuropathogenesis. Taking advantage of its flexibility for genetic manipulations, this innovative model provides an unprecedented access to the infected brain and “hard-to-catch” cell populations to study ZIKV pathogenesis as well as to perform *in vivo* drug screening and testing.

## METHODS

### Fish husbandry

Adult zebrafish (*Danio rerio*) were reared at 28°C with a 12/12 light-dark cycle in the aquatic facility of the National Laboratory of Experimental Biology at Institut National de la Recherche Scientifique (INRS). Fertilized eggs were collected, treated as specified, kept in petri dishes, and raised at 28.5°C. The zebrafish lines used were: wild-type TL, Tg(*nestin*:GFAP), Tg(*gfap*:GFP) and Tg(*dlx5a/6a*:GFP;*vglut2*:RFP). The *nestin* and *gfap* transgenic lines, previously generated and described^45,46^, were obtained from the laboratories of Pierre Drapeau and Eric Samarut. Tg(*dlx5a/6a*:GFP) was obtained from Dr. Marc Ekker. All the experiments were performed in compliance with the guidelines of the Canadian Council for Animal Care and under the approval of local ethic and biosafety committee of INRS.

### ZIKV and DENV production

Zika virus strain H/PF/2013 was obtained from European Virus Archive Global (EVAg) and was passaged in Vero E6 cells.^84^ The plasmid coding for DENV2 16681s genome sequence (pFK-DENV-WT) was previously described.^85^ Following pFK-DENT-WT linearization with XbaI, *in vitro* transcription was performed using mMessage mMachine kit (Thermo-Fisher) with SP6 RNA polymerase. DENV2 16681s stock were produced after electroporation of *in vitro* transcribed genomes in Vero E6 cells as described in ^86^. ZIKV and DENV viral titers were determined by plaque assay. Briefly, 24-well plates were seeded with 2 x 10^5^ Vero E6 cells/well. Cells were infected with serially diluted virus samples. Two hours post-infection, inoculums were replaced for MEM (Invitrogen) containing 1.5% carboxymethylcellulose (Millipore-Sigma). 5 days (ZIKV) and 7 days (DENV) after infection, cells were fixed with 10% formaldehyde and plaques were counted following staining with 1% crystal violet/10% ethanol.

### Embryo infection

Using the FemtoJet 4i microinjector (Eppendorf), a volume of 2 nL containing ZIKV particles (∼5 x 10^7^ PFU/mL), DENV particles (∼4 x 10^6^ PFU/mL) or DMEM (vehicle) was injected into 2-to-4 cell stage embryos. Four hours post-injection, fish were either treated with 100 μM NITD008 (Tocris Small Molecules) or 0.5% DMSO as control.

### Viral protein expression in embryos

The plasmids containing NS4A, 2k-NS4B, or NS4A-NS4B sequences downstream T7 RNA polymerase promoter were previously described.^87^ Following pTM-ZIKV NS4A, pTM-ZIKV 2K-NS4B, and pTM-ZIKV NS4A-NS4B plasmids linearization with SpeI, *in vitro* transcription was performed using the TranscriptAid T7 High Yield transcription kit (Thermo Scientific). RNA translation initiation is driven by the encephalomyocarditis virus (ECMV) internal ribosome entry site (IRES) at the 5’ terminus. A volume of 1 nL containing 500 ng/μL mRNA coding for each of these proteins was injected into one-cell stage embryos using the FemtoJet 4i microinjector (Eppendorf). H_2_O was used as reference control.

### Survival and morphology

Injected embryos were monitored for their survival rate for 3 days post-fertilization using heartbeat as a readout of viability. At the latest time point (*i.e.*, 3 days post injection larval stage), larvae were observed under Stemi 305 microscope (Zeiss) to assess hatching and general morphology. For head area measurement, larvae were positioned laterally, and head sizes were calculated using the Fiji software.

### Behavioral analysis

At 4 days post-fertilization, larvae were separated into single wells of a 96-wells plate containing 200 μL of E3 media. Following 30 minutes of habituation in the Daniovision recording chamber (Noldus), locomotor activity upon light stimulation was monitored for two hours. Swimming distance (in mm) and immobility times (in seconds) analysis were performed using the EthoVision XT12 software (Noldus). Swimming and responses-to-touch videos were taken with an iPhone (Apple) coupled to a stereomicroscope at 30 frames per second.

### RT-qPCR and RT-ddPCR

At the indicated time points, RNAs were isolated from whole infected or non-infected larvae using the RNeasy mini kit (Qiagen) according to the manufacturer’s protocol. Extracted RNAs were subjected to reverse transcription using the Invitrogen SuperScript IV VILO Master Mix RT kit (Life Technologies) or QuantiTect Reverse Transcription kit (Qiagen). ZIKV RNA and *vglut1* mRNA were detected by droplet digital PCR (ddPCR) and absolute quantification was performed using the QX200 ddPCR Evagreen Supermix (Bio-Rad) and the QX200 ddPCR System (Bio-Rad). For other RNA quantifications, real-time PCR was performed using the Applied Biosystems SYBR Green Master mix (Life Technologies) and a LightCycler 96 (Roche) for the detection. The following primer pairs were used: 5′-AGATGAACTGATGGCCGGGC*-*3′ and 5′*-* AGGTCCCTTCTGTGGAAATA*-*3′ for ZIKV H/PF/2013; 5’-GCCCAAGTAAACACCCTGGA-3’ and 5’-GCAAAGCTCTGTATGTTGCCA-3’ for *nestin*; 5’-AATGTCAAACTGGCCCTGGA-3’ and 5’-TCTGCACCGGAACAGTGATT-3’ for *gfap*; 5’-CTTTAAAGCCCAGGCAGGGA-3’ and 5’-AGGATGGCGATGATGTAGCG-3’ for *vglut1*; 5’-TGAGAACTCAAATGTGGACCT-3’ and 5’-GTCCTCCACCTTTGACTTGT-3’ for *ifnφ1*; 5’-CGGACCTCAGTTTCAAGG-3’ and 5’-GCAGCGGGAGAATATGGA-3’ for *rig-I*; 5’-GTGGCTGGAGACAGCAAGA-*3’* and *5’*-AGAGATCTGACCAGGGTGGTT-*3’* for *ef1a*. The ΔΔCt method was used to determine the relative expression levels normalized to *ef1a*, used as a housekeeping gene.

### TUNEL assays

To analyze cell death, TUNEL assays were performed on whole embryos at 2 days post-fertilization. At one day post-fertilization, embryos were treated with 0.003% phenyltheioura (PTU) to block pigment formation. One day later (*i.e.*, two days post injection), larvae were fixed overnight with 4% paraformaldehyde (PFA, in PBS), dehydrated, and rehydrated using serial dilutions of methanol in PBST (PBS containing 0.1%Tween) then washed with PBST. Following digestion with proteinase K at 10 μg/mL for 20 minutes, larvae were washed with PBST and PBS, fixed with 4% PFA/PBS for 20 minutes, and incubated with the *In Situ* Cell Death Detection Kit Fluorescein mix (Roche) at 37°C for 1 hour. Larvae were imaged with a LSM780 confocal microscope (Carl Zeiss Microimaging) at the Confocal Microscopy and Flow Cytometry Core Facility of INRS.

### Brain cryo-sections and immunofluorescence

At 2 days and 4 days post-fertilization, zebrafish were fixed in 4% PFA/PBS, dehydrated in serial dilutions of sucrose, and frozen in Tissue plus O.C.T compound (Fisher Scientific). 12 micron-thick transverse sections of the head were prepared, dried at room temperature and frozen at −80°C until use.

For Sox2 staining and Sox2/ZIKV E co-staining, sections were washed with PBS. Antigen retrieval was performed solely on Sox2 single staining by incubation with Tris-HCl (pH 8.2, 50mM) at 85°C for 6 minutes. Sections were then washed with PBS-0.5% TritonX-100, incubated in blocking solution [10% normal goat serum (NGS) in PBS] for one hour, and in primary antibody solution anti-Sox2 (1:200, Invitrogen), panflaviviral anti-E (clone 4G2; 1:400, Genetex) diluted in 5% NGS, 1% bovine serum albumin in PBS-0.1%TritonX-100 overnight at 4°C. After several washes with PBS-0.3% TritonX-100, sections were incubated with species-specific Alexa Fluor-conjugated secondary antibodies (Life Technologies) for 2 hours at room temperature in the dark. Sections were washed for one hour with PBS-0.3% TritonX-100, incubated for 10 minutes in 1:1000 Hoescht to stain the nuclei, and mounted.

Transversal brain sections were imaged with a LSM780 confocal microscope (Carl Zeiss Microimaging) and were analyzed using Zeiss Zen (black edition) and Fiji software. Schematic brain representation and structures’ identification were made based on the Atlas of Early Zebrafish Brain Development by Mueller and Wulliman ^51^.

### Embryos dissociation and flow cytometry

Tg(*gfap*:GFP) and Tg(*nestin*:GFP) embryos dissociation was performed as described before ^48^ with minor modifications. One day post-fertilization embryos were dechorionated, and deyolked in deyolking buffer (55 mM NaCl, 1.8 mM KCl, 1.25 mM NaHCO_3_) in the presence of 10,000 123count eBeads counting beads (Thermo-Fisher). Deyolked embryos were pelleted at 500g for 5 minutes and rinsed with FACSmax cell dissociation solution (Genlantis). Embryos were transferred in 6 well plates and dissociated with FACSmax by gentle up-and-downs. Single cells suspensions were pelleted and washed with PBS. For flow cytometry experiments, cells were fixed in 4% PFA/PBS and filtered using a cell strainer (Fisher Scientific) before data acquisition using a LSRFortessa Cell Analyzer (BD) at the Confocal Microscopy and Flow Cytometry Core Facility of INRS. Data analysis was performed using the FlowJo software (version 10.8.1).

### Cell sorting and RNA-sequencing

One day post injection Tg(*nestin*:GFP) embryos were dissociated into single-cells suspension as described above. Cell sorting was performed on unfixed cells using a FACSMelody Cell Sorter (BD Biosciences) in a biosafety level 2 cabinet located at the Confocal Microscopy and Flow Cytometry Core Facility of INRS. GFP expressing cells were directly sorted into buffer RLT (Qiagen) containing β-mercaptoethanol. RNAs were extracted using the RNeasy Mini kit (Qiagen) according to the manufacturer’s instructions. RNA quality control and next generation RNA sequencing were performed at the Genomics Core Facility of the Institute for Research in Immunology and Cancer (IRIC, University of Montreal) using 2100 bioanalyzer (Agilent) and Illumina NextSeq 500 instrument, respectively. RNA sequencing data analysis was performed by the Bioinformatics Core Facility of IRIC.

### Statistical analyses

Data were analyzed using GraphPad Prism 8 software. The number of experiments (N) and sample size (n) are mentioned in the legends. Dare are presented as mean ± SEM or median ± 95% confidence interval (CI) as specified in legends. Normality test was determined using the D’Agostino & Pearson test. Significance was determined using either unpaired Student’s t-test, one-way ANOVA, two-way ANOVA (for data presenting a normal distribution), nonparametric Mann-Whitney or nonparametric Kruskal-Wallis (for data presenting a non-normal distribution) tests as specified in legends.

## ACKNOWLEDGEMENTS

The authors would like to thank Charlotte Zaouter for her valuable help in zebrafish care and maintenance. We are grateful to Jessy Tremblay at the Confocal Microscopy and Flow Cytometry Facility of the Armand-Frappier Santé Biotechnologie research center for excellent technical assistance during imaging and cell sorting. For help with sequencing data analysis, we thank Patrick Gendron at Bioinformatics Core Facility of the Institute for Research in Immunology and Cancer (University of Montreal). We are grateful to the European Virus Archive Global (EVAg) and Dr. Xavier de Lamballerie (Emergence des Pathologies Virales, Aix-Marseille University, France) for providing ZIKV original stocks. We thank Dr. Pei-Yong Shi and the World Reference Center for Emerging Viruses and Arboviruses (WRCEVA), and Dr. Ralf Bartenschlager (University of Heidelberg) for providing the ZIKV and DENV molecular clones. A.A.S has received PhD fellowships from the Fonds de la Recherche du Québec-Nature et Technologies (FRQNT), the Armand-Frappier Foundation and the Center of Excellence in Research on Orphan Diseases-Courtois Foundation (CERMO-FC). P.J was supported by a CERMO-FC scholarship. L.C.C is receiving a research scholar (Senior) salary support from FRQS. SP holds an FRQS Junior 2 research scholar award and the Anna Sforza Djoukhadjian Research Chair. This study was supported by the Canada Foundation for Innovation (CFI; #37512, to S.P), by the Canadian Institutes for Health Research (CIHR; PJT 190064 to L.C.C and S.P, and PJT 177940 to S.P), by an Acceleration grant from CERMO-FC to S.P and L.C.C, and by a Azrieli Future Leader in Canadian Brain Research grant from Brain Canada through the Canada Brain Research Fund, with the financial support of Health Canada and the Azrieli Foundation awarded to L.C.C.

---

Figure S1. Interferon response and mobility following ZIKV injection.

(A) At 3 days post-fertilization, mRNA amounts of *rig-I* and ifnφ1 in mock and ZIK-infected larvae were analyzed by RT-qPCR. N=4.

(B and C) The distance moved (B) by the larvae and the immobility time (C) were assessed using the DanioVision device (mock; *n*=27; ZIKV, *n*=35; DENV, *n*=34. *N*=2). Data are shown as median ± 95 CI. **** P ≤ 0.0001; *** P ≤ 0.001; ** P ≤ 0.01; Kruskal-Wallis test.

*n* indicates the number of fish; *N* represents the number of independent experimental repeats.

Figure S2. ZIKV infection of Tg(*gfap*:GFP) and Tg(*nestin*:GFP) induces morphological defects and a depletion of NPCs.

(A) Proportion of Tg(*gfap*:GFP) larvae with different phenotypes at 3 days post-fertilization following ZIKV-injection (*n*=27) or mock-injection (*n*=32). *N*=1.

(B) Proportion of Tg(*nestin*:GFP) larvae with different phenotypes at 3 days post-fertilization following ZIKV-injection (*n*=85) or mock-injection (*n*=87). *N*=2.

(C) At 1 day post-fertilization, ZIKV infected or uninfected whole transgenic Tg(*nestin*:GFP) embryos were dissociated in the presence of 10,000 fluorescent normalizing beads. Single cells and beads were counted by flow cytrometry. Number of GFP^+^ cells counted per 100 beads are shown (N=1).

(D and E) At 1 day post-fertilization, mRNA amounts of *nestin* and *gfap* in (GFP^+^ (NPCs) and GFP-in mock and ZIK-infected tg(*gfap*:GFP) (D) and tg (*nestin*:GFP) embryos (E) were analyzed by RT-qPCR.

*n* indicates the number of fish; *N* represents the number of independent experimental repeats.

Figure S3. ZIKV infection induces a loss of NPCs in larvae at 4 days post-fertilization.

(A) Schematic representation of a zebrafish brain at 4 days post-fertilization. The three areas of the brain are shown: forebrain, midbrain, and forebrain. Gray lines represent the localization of the transverse sections. D= dorsal; V= ventral; A= anterior; P= posterior.

(B-D) Number of neural progenitor cells (Sox2^+^ cells) in the midbrain (B) and the hindbrain (C) of 4 dpf mock-injected or ZIKV-injected fish. TeO= tectum opticum; T= midbrain tegmentum; N= region of the nucleus of medial longitudinal fascicle; MO= medulla oblongata. Scale bars = 50 μm.

(D) Quantification of (B-C). Data are means ± SEM. ** P ≤ 0.01; * P ≤ 0.05; ns: non-significant Two-way ANOVA.

Figure S4. Injection of the ZIKV RNA genome into embryos induces morphological defects.

(A) *In vitro* transcribed ZIKV RNA genome (vRNA) was microinjected in the zebrafish embryo at 1 hour post-fertilization.

(B) Representative pictures of microinjected larvae at 3 days post-fertilization. ZIKV vRNA injection induced both severe and mild developmental phenotypes.

(C and D) Quantification of the proportion of larvae with the different phenotypes (C), and head size (D) at 3dpf of the larvae (Mock, *n*=19; ZIKV RNA, *n*=22. *N*=2). Data are are means ± SEM. * P ≤ 0.05; T-test. *n* indicates the number of fish; *N* represents the number of experimental repeats.

**Video S1:** Locomotor activity of control (mock), ZIKV-infected (ZIKV), and ZIKV-infected and NITD008-treated (ZIKV+NITD008) embryos at 2 days post-fertilization.

**Video S2:** Locomotor activity of control (mock), ZIKV-infected (ZIKV), and ZIKV-infected and NITD008-treated (ZIKV+NITD008) larvae at 3 days post-fertilization.

**Video S3:** Locomotor activity of water control (mock) and ZIKV NS4A-encoding RNA-injected larvae at 3 days post-fertilization.

**Table S1:** RNA-seq differential expression analysis of NPCs isolated from ZIKV-infected and control embryos.

